# VAV2-Dependent Regulation of Ribosome Biogenesis in Keratinocytes and Oral Squamous Cell Carcinoma

**DOI:** 10.1101/2023.11.03.565464

**Authors:** Natalia Fernández-Parejo, L. Francisco Lorenzo-Martín, Juana M. García-Pedrero, Juan P. Rodrigo, Mercedes Dosil, Xosé R. Bustelo

## Abstract

VAV2 is an activator of RHO GTPases that promotes and maintains regenerative proliferation-like states in normal keratinocytes and oral squamous cell carcinoma (oSCC) cells, respectively. Here, we demonstrate that VAV2 also plays critical roles in the regulation of ribosome biogenesis in those cells, a program associated with poor prognosis of human papilloma virus negative oSCC patients. Mechanistic analyses indicate that VAV2 regulates this process in a catalysis-dependent manner using a conserved pathway composed of the GTPases RAC1 and RHOA, members of the PAK and ROCK family kinases, and the transcriptional factors c-MYC and YAP/TAZ. This pathway directly promotes RNA polymerase I activity and the ensuing synthesis of 47S pre-rRNA precursors. This process is further consolidated by the upregulation of ribosome biogenesis factors and the acquisition of the YAP/TAZ-dependent cell undifferentiation state. Finally, we show that RNA polymerase I is a therapeutic Achilles’ heel for both keratinocytes and hnSCC patient derived cells endowed with high VAV2 catalytic activity. Collectively, these findings highlight the therapeutic potential of the VAV2 and ribosome biogenesis pathways in both preneoplastic and late progression stages of oSCC.

## INTRODUCTION

Head and neck squamous cell carcinoma (hnSCC) can develop in the epithelia of the mucosal lining of the upper aerodigestive tract areas such as the oral epithelium, the tongue, the larynx, and the hypopharynx. These tumors are clinically challenging due to epidemiological incidence, metastatic properties, frequent posttreatment recurrence events, and limited therapeutic options. Factors influencing the development of these tumors include alcohol intake, tobacco smoking, and human papilloma virus (HPV) infections (1). Approximately 670 000 hnSCC cases have been detected worldwide in 2020, which show an average 40 to 50% mortality rate (2).

Recent work has revealed a number of biological traits that favor hnSCC development and malignant properties (1, 3). One of them is regenerative proliferation, a biological process characterized by the presence of high percentages of proliferating and undifferentiated cells that correlates with poor hnSCC prognosis (1, 4). This trait can be orchestrated in a concerted manner by multiple transcriptional factors such as the YAP/TAZ complex, AP1, E2F, c-MYC, TP63, and ACTL6A (5–10). More recently, it has been shown that VAV2, a tyrosine phosphorylation-regulated guanosine nucleotide exchange factor (GEF) that catalyzes the activation step of the GTPases RAC1 and RHOA (19–22), also plays critical roles in early signaling events that trigger and maintain regenerative proliferation in normal keratinocytes and oSCC, respectively (18). VAV2 regulates this process through the engagement of the GTPases RAC1 and RHOA, the stimulation of the proximal GTPase effectors PAK and ROCK, and the activation of the transcriptional factors c-MYC and YAP/TAZ. These two factors are responsible for high proliferative (c-MYC) and undifferentiation state (YAP/TAZ) of cells (18). The levels of VAV2 mRNA levels and VAV2-regulated gene signatures directly correlate with poor HPV^−^ hnSCC patient prognosis (18), further underscoring the importance of this pathway for the malignant properties of this tumor type. In line with this, it has been shown using xenotransplant experiments that the knockdown of endogenous *VAV2* reduces the primary tumorigenesis and metastatic properties of oSCC patient derived cells (PDCs) (18).

Recent evidence indicates that hnSCCs also depend on high ribosome biogenesis rates for optimal fitness (11). This biological process is initiated by the RNA polymerase I-mediated transcription of the 47S pre-ribosome RNA (rRNA) in the nucleolus. This precursor subsequently undergoes several cleavage and maturation steps in the nucleolus, nucleoplasm and cytosol that lead to the generation of the rRNAs that form part of both the small (18S rRNA) and large (28S rRNA, 5.8S rRNA) ribosome subunits. This maturation process is accompanied by the sequential docking and release of ribosome biogenesis factors and the final incorporation of ribosomal proteins (12, 13). This process is targeted by many oncogenic drivers and signal transduction pathways in cancer cells (12). Connected to this route, it been shown that the translation of specific transcript subsets via deregulation of translational regulators also contributes to hnSCC fitness (14–16).

Despite this progress, very little information is still available regarding the potential interconnections established by all the foregoing biological and signaling pathways in hnSCC. For example, despite the extensive functional characterization of RHO proteins at the signaling and cellular level during the last decades, the influence of these GTPases on ribosome biogenesis remains ill characterized in hnSCC and other tumor types. Recent observations indicate that such interconnections might indeed exist, although they entail rather noncanonical and tumor type specific signaling mechanisms. For example, in the case of non-small cell lung cancer, it has unexpectedly been found that the RHO GEF ECT2 and RAC1 promote direct RNA polymerase I transcription through the interaction with the nucleolar protein nucleophosmin (17). Conversely, a negative regulator of RHO GTPases (ARHGAP30) has been shown to negatively regulate ribogenesis in cervical cancer. This mechanism relies on the ubiquinylation-mediated degradation of a key ribosome biogenesis factor rather than on the expected regulation of RHO GTPase activity (18). To date, however, we do not know whether these or other alternative mechanisms operate in hnSCC. Potential interconnections between ribogenesis and regenerative proliferation also remains poorly characterized.

Using organotypic cultures of primary keratinocytes and oSCC PDCs as experimental model, we here present evidence demonstrating that VAV2 coordinates the concurrent regulation of regenerative proliferation and ribogenesis a RAC1 and RHOA GTPase-dependent manner in both normal and fully transformed cells. This connection is therapeutically relevant, since the VAV2-regulated gene signature for ribosome biogenesis factors is associated with poor prognosis of HPV^−^ oSCC patients. Perhaps more importantly, we show that ribogenesis represents a key therapeutic vulnerability for keratinocytes and oSCC PDCs exhibiting high levels of VAV2 activity.

## RESULTS

### VAV2^Onc^-driven epidermal hyperplasia correlates with enhanced ribogenesis

We previously found using *in silico* annotation analyses that ribosome ribogenesis was one of top upregulated biological functions in the hyperplasic epidermis of *Vav2*^Onc/Onc^ mice (19). These knock-in mice endogenously express a truncated version of VAV2 (Δ1-186, referred to hereafter as VAV2^Onc^) that shows constitutive catalytic activity due to the removal of the two autoinhibitory N-terminal domains (19). As a result, this leads to the chronic stimulation of the downstream RAC1 and RHOA GTPases (19). In line with those *in silico* analyses, we found using gene set enrichment analyses (GSEA) that gene signatures for both ribosome biogenesis factors (**Supplementary** Fig. 1A and B) and structural ribosomal proteins (**Supplementary** Fig. 1C and B) are highly enriched in the VAV2^Onc^-upregulated transcriptome. Further analyses indicated that the expression levels of the VAV2^Onc^-regulated transcripts for ribosome biogenesis factors are also increased in oSCC when compared to either healthy or dysplastic tissue samples (**Supplementary** Fig. 1D, top panel). The expression levels of the subset of VAV2^Onc^-regulated ribosome biogenesis factor-encoding transcripts also correlate with the abundance of the *VAV2* mRNA (**Supplementary Fig. S1E**, top panel) and with poor patient prognosis (**Supplementary Fig. S1F**, top panel) when tested in a cohort of HPV^−^ oSCC patients (20). The stratification power of this signature (*P* = 0.016) is similar to that provided by the expression of the *EGFR* transcript (*P* = 0.011), a gene with key protumorigenic functions in hnSCC (1, 3). However, its power is significantly lower than the provided by the expression levels of the *VAV2* mRNA itself (*P* = 0.008) (19) as well as of other raw (*P* = 0.009) and refined (*P* < 1 x 10^−5^) VAV2^Onc^-regulated gene signatures (19). The ribosome biogenesis gene signature cannot stratify patients well when using gene expression datasets lacking information on HPV status (L.F.L.-M. and X.R.B., data not shown), suggesting that its functional relevance is limited to HPV^−^ hnSCC cases. We could not observe any correlation of the levels of the gene signature for structural components of mature ribosomes with tumor progression (**Supplementary** Fig. 1D, bottom panel), *VAV2* mRNA abundance (**Supplementary** Fig. 1D, bottom panel), or patient prognosis (**Supplementary** Fig. 1C, bottom panel) using the foregoing gene expression datasets. These analyses suggest that the upregulation of ribogenesis might play pathogenic roles in VAV2^Onc^-dependent HPV^−^ oSCC subtypes.

Given the foregoing results, we decided to investigate whether ribosome biogenesis was upregulated in VAV2^Onc^-expressing primary keratinocytes and, if so, whether it contributed to the epithelial hyperplasia induced by this constitutively active protein. To this end, we utilized as main working model organotypic three dimensional (3D) cultures of primary mouse or human keratinocytes. This strategy allowed us to monitor the regulation of ribosome biogenesis in a tissue-like model that recapitulates all the differentiation stages of keratinocytes (19). When using keratinocytes from newborn wild-type (WT) and *Vav2*^Onc/Onc^ knock-in mice, we found as previously reported (19) that the cells with endogenous expression of VAV2^Onc^ generate thicker layers of suprabasal cells than the WT counterparts (**Fig. 1A**). This is due to the proliferative expansion of highly undifferentiated keratinocytes located in the suprabasal layer (19). We stained sections from those 3D cultures with an antibody to the 5.8S rRNA (5.8S), an integral component of the large 60S ribosome subunit that is generated from the 47S pre-rRNA precursor (12), to assess ribogenesis activity in all cell layers of the epithelia formed. Using this approach, we found that the epithelia generated by WT keratinocytes show high levels of ribogenesis in the basal layer and, to a much lesser extent in suprabasal cells (**Fig. 1A**). By contrast, the hyperplastic epithelia formed by *Vav2*^Onc/Onc^ keratinocytes display high levels of 5.8S rRNA immunoreactivity in both basal cells and in the highly expanded layer of suprabasal cells (**Fig. 1A** and **B**).

**FIGURE 1.**
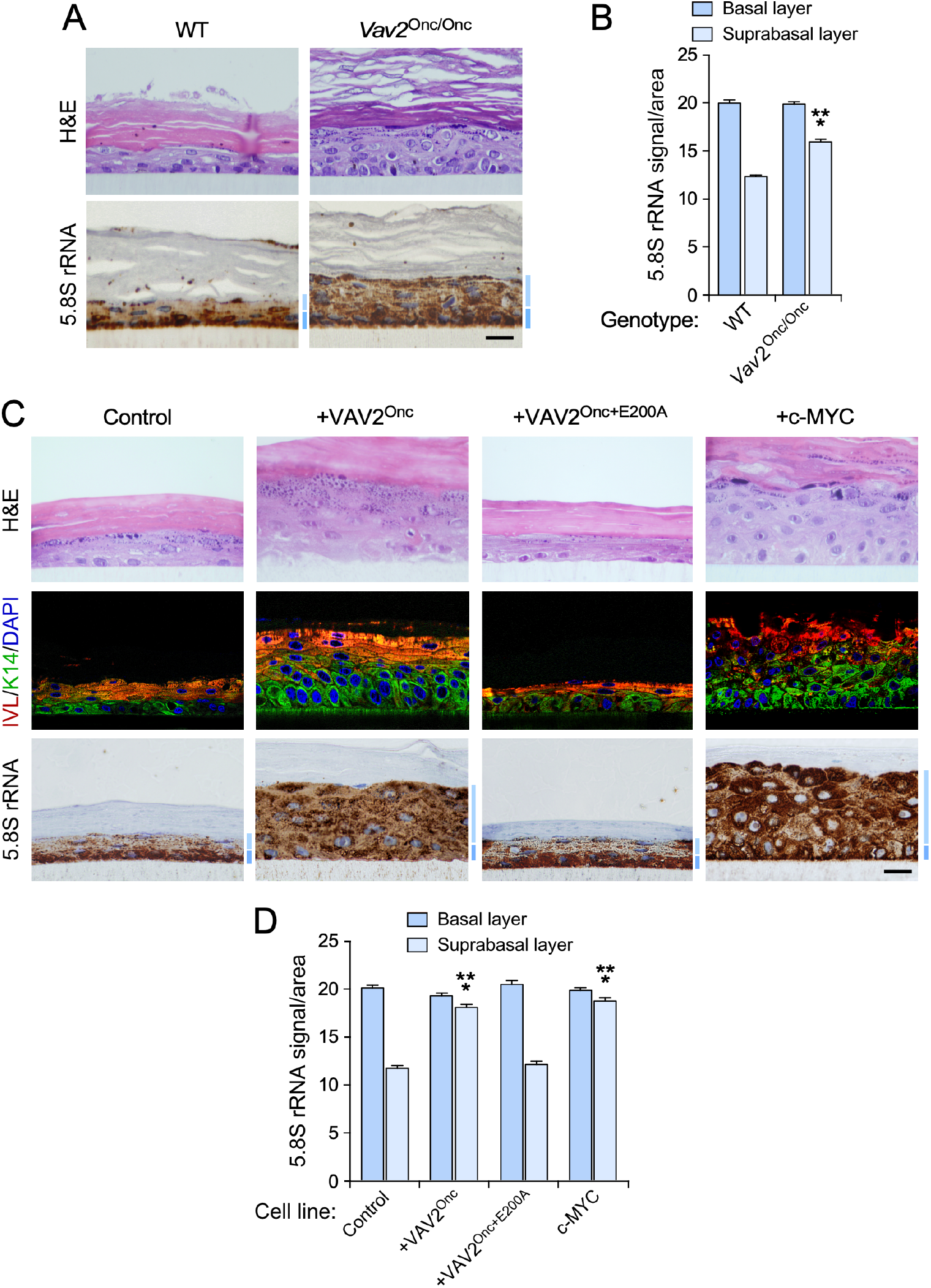
VAV2^Onc^-driven epidermal hyperplasia correlates with enhanced ribogenesis. (**A**) Histological sections from organotypic cultures using primary keratinocytes from WT and *Vav2*^Onc/Onc^ mice that were stained with either hematoxylin-eosin (H&E, top) or antibodies to the 5.8S rRNA antibody plus hematoxylin. The basal and suprabasal epidermal layers are indicated by a dark blue and light blue bar (bottom panel), respectively. Scale bar, 10 μm. **(B)** Quantification of the 5.8S rRNA immunoreactivity obtained in the basal and suprabasal layers from (A). ***, *P* < 0.0001 (Student’s *t*-test, *n* = 3 independent cultures). **(C)** Tissue sections of 3D organotypic models using human keratinocytes expressing the indicated proteins upon staining with either hematoxylin-eosin (top panels) or antibodies to involucrin (IVL, red color), keratin 14 (K14, green color, middle panels), and the 5.8S rRNA (brown color, bottom panels). Some of the sections were subsequently counterstained with either 4′,6-diamidino-2-phenylindole dihydrochloride (DAPI, blue color, middle panels) or hematoxylin (bottom panels). Dark and light blue bars indicate the basal and suprabasal layers, respectively. Scale bar, 10 μm. **(D)** Quantification of the 5.8S rRNA immunoreactivity in the organotypic cultures displayed in (C). ***, *P* < 0.0001 (ANOVA and Dunnett’s multiple comparison test, *n* = 5 independent cultures. In (B) and (D), data represent the mean ± SEM. Source data for this figure are provided as a Source Data file.

We next carried out similar analyses using primary human keratinocytes stably expressing catalytically active (VAV2^Onc^) or catalytically inactive (VAV2^Onc+E200A^) versions of VAV2. VAV2^Onc+E200A^ contains a point mutation in the catalytic DBL-homology domain that eliminates the enzyme activity of the protein (19). As negative control, we used primary human keratinocytes expressing an empty lentivirus. As positive control we used cell derivatives ectopically expressing c-MYC, a transcriptional factor known to be involved in ribosome biogenesis (12). The generation, validation, and biological characterization of all these keratinocyte lines has been reported before (19). As previously described (19), we observed that keratinocytes expressing either VAV2^Onc^ or c-MYC promoted exacerbated levels of epithelial hyperplasia when tested in organotypic 3D cultures (**Fig. 1C**, upper panels). As in the case of Vav2^Onc/Onc^ keratinocytes, this hyperplasia is the result of the expansion of highly proliferative and undifferentiated cells located in the suprabasal layer (**Fig. 1C**, middle panels) (19). By contrast, keratinocytes expressing VAV2^Onc+E200A^ generated control-like epithelial structures under the same experimental conditions (**Fig. 1C**, upper and middle panels) (19). This is consistent with the fact that the VAV2^Onc^-driven hyperplasia relies on a RHO GTPase–c-MYC signaling axis (19). The immunostaining of those sections with antibodies to the 5.8S rRNA revealed increased levels of this rRNA in the expanded layers of suprabasal cells formed in the organotypic cultures of both VAV2^Onc^– and c-MYC-expressing keratinocytes (**Fig. 1C** and **D**). The stable expression of VAV2^Onc+E200A^ in keratinocytes does not change the usual pattern of 5.8S rRNA immunoreactivity found in the epithelial structures formed by control cells (**Fig. 1C** and **D**), further indicating that the effects elicited by VAV2^Onc^ on the distribution of 5.8S rRNA immunoreactivity are catalysis dependent. Consistent with this idea, we demonstrated using similar organotypic 3D culture experiments that the stable expression of constitutively active versions of RAC1 (F28L mutant), RHOA (F30L mutant), CDC42 (F28L mutant) or the combination of RAC1^F28L^ plus RHOA^F30L^ promotes a redistribution of 5.8S rRNA immunoreactivity in the organotypic sections very similar to that seen in VAV2^Onc^– and c-MYC-expressing keratinocytes (**Supplementary** Fig. 2). All these GTPase mutant versions become chronically active owing to accelerated GDP/GTP exchange rates (21). In line with previous results (19), we found that the hyperplasic layer was significantly thicker in the organotypic cultures generated using RAC1^F28L^+RHOA^F30L^-expressing human keratinocytes (**Supplementary** Fig. 2).

As an additional read-out to confirm the increased ribogenic activity in VAV2^Onc^-expressing keratinocytes, we next analyzed the number and size of the nucleoli present in our collection of human keratinocyte derivatives. Nucleoli are the nuclear structures where rRNA is transcribed and initially processed, so their structure can undergo significant changes depending on the ribogenic activity of cells (12). When compared to controls, we found that VAV2^Onc^-, c-MYC-, and RAC1^F28L^+RHOA^F30L^-expressing keratinocytes show much larger nucleoli than controls (**Supplementary** Fig. 3A and B). This is connected with a reduction in the average number of nucleoli present per cell. (**Supplementary** Fig. 3A and C). Taken together, these results indicate that the constitutive activation of the catalytic-dependent pathways of VAV2 promotes enhanced ribogenesis in both mouse and human keratinocytes.

### VAV2^Onc^ promotes rRNA synthesis in an RNA polymerase I-dependent manner

To investigate how VAV2^Onc^ promotes enhanced levels of ribosome biogenesis, we first analyzed its impact on the synthesis of the primary 47S pre-rRNA precursor. To this end, we used pulse chase experiments with 5-ethynil uridine (5-EU) followed by click chemistry-based reactions with an Alexa Fluor™ 594 carboxamido-(6-azidohexanyl), triethylammonium salt to label the nascent 47S pre-RNA precursors in exponentially growing 2D cultures of control, VAV2^Onc^-, VAV2^Onc+E200A^-, and c-MYC-expressing cells. We found that VAV2^Onc^-expressing cells and, to a larger extent c-MYC-expressing keratinocytes display higher levels of 5-EU incorporation than controls (**Fig. 2A**, **B** and **C**). By contrast, the keratinocytes expressing the catalytically dead version of VAV2 display levels of 5-EU incorporation similar to controls (**Fig. 2A**, **B** and **C**). The addition of an RNA polymerase I inhibitor (CX-5461) to the cultures brings the levels of 5-EU incorporation of VAV2^Onc^– and c-MYC-expressing cells back down to those observed in control cells (**Fig. 2C**). These results suggest that the elevated levels of 47S pre-RNA production in VAV2^Onc^– and c-MYC-expressing keratinocytes is due to the activation of that polymerase. This is probably the result of direct signaling, since we observed using luciferase reporter assays that VAV2^Onc^ can also promote RNA polymerase I activity when transiently transfected in primary human keratinocytes using 2D cultures (**Fig. 2D** and **E**).

**FIGURE 2.**
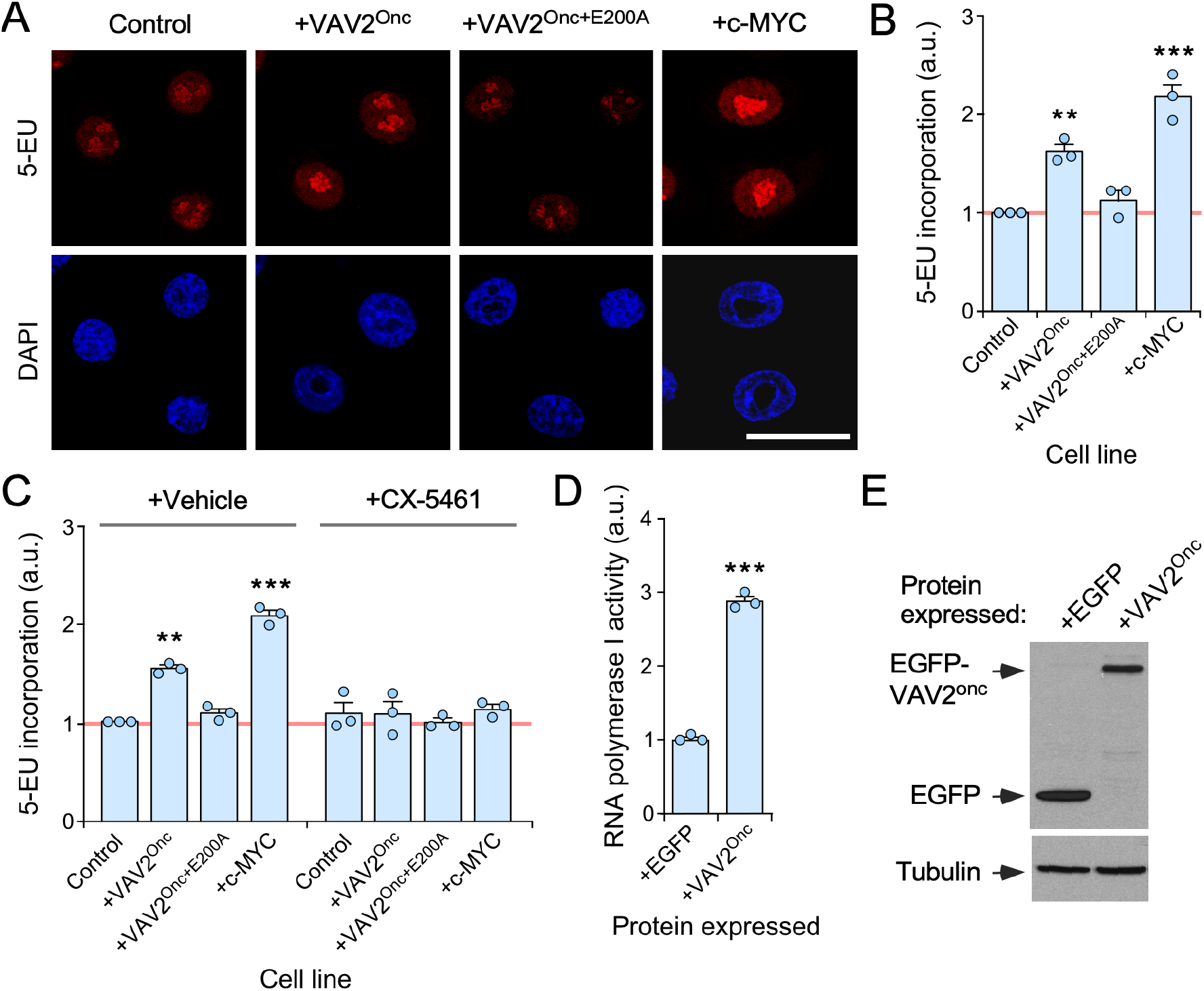
VAV2^Onc^-driven epidermal hyperplasia is associated with enhanced ribogenesis. (**A**) Representative images of 5-EU– (red color, top panels) and DAPI-labeled (blue color, bottom panel) human keratinocytes expressing the indicated proteins (top). Scale bar, 20 μm. **(B)** Quantitation of the incorporation of 5-EU found in the experiments shown in (A). **, *P* = 0.003; ***, *P* < 0.0001 of indicated test samples versus control cells (ANOVA and Dunnett’s multiple comparison test, *n* = 3 independent experiments). a.u., arbitrary units. **(C)** Quantitation of the incorporation of 5-EU in indicated keratinocyte lines upon inhibition of RNA polymerase I with CX-5461. **, *P* = 0.006; ***, *P* < 0.0001 (experimental test vs controls) (ANOVA and Tukey’s HSD tests, *n* = 3 independent experiments). **(D)** Determination of RNA polymerase I activity of keratinocytes transiently expressing the indicated proteins using a luciferase reporter assay. Values have been normalized to the experimental value obtained in EGFP-expressing cells (which was given an arbitrary value of 1). *, *P* = 0.017; ***, *P* < 0.0001 (ANOVA and Tukey’s HSD tests, *n* = 3 biological replicates). **(E)** Immunoblot showing the expression of the ectopically expressed proteins in one of the experiments performed in (D) (top panel). Tubulin α was used as protein loading control (bottom panel). Same results were obtained in two independent experiments (not shown). In (B) to (D), data represent the mean ± SEM. Source data for this figure are provided as a Source Data file.

We did not find any statistically significant change in the abundance of pre-rRNA intermediaries that participate in the maturation stages of either the small (18S rRNA containing) or the large (28S rRNA containing) ribosome subunits in any of the interrogated keratinocyte lines (**Supplementary** Fig. 4). These findings indicate that VAV2^Onc^ primarily affects the RNA polymerase I-mediated synthesis of 47S pre-rRNA precursors in a catalytically dependent manner.

### Mechanistic analysis of VAV2^Onc^-driven ribogenesis

We used the 5-EU labeling method to obtain further mechanistic information on how VAV2^Onc^ promotes ribosome biogenesis in keratinocytes. Consistent with being a VAV2 catalysis-mediated mechanism (see above, **Figs. 1** and **2**), we found that keratinocytes ectopically expressing active RAC1^F28L^ plus RHOA^F30L^ show increased rates of 5-EU incorporation similar to those induced by VAV2^Onc^ (**Fig. 3A** and **B**). Next, we utilized inhibitors for downstream elements of the VAV2^Onc^ pathway that were previously shown to be important for maintaining the regenerative proliferation in both mouse and human primary keratinocytes (19) (**Fig. 3C**). Those included compounds that target RAC1 (1-A116), PAK (FRAX597), ROCK (Y27632), c-MYC (10058-F4), and the YAP/TAZ complex (verteporfin) (**Fig. 3C**). All these inhibitors eliminated the high levels of 5-EU-labeled precursors when added to VAV2^Onc^-expressing cells (**Fig. 3D** to **F**). However, none of them elicited any statistically significant change in the basal levels of 5-EU incorporation found in control cells (**Fig. 3D** to **F**).

**FIGURE 3.**
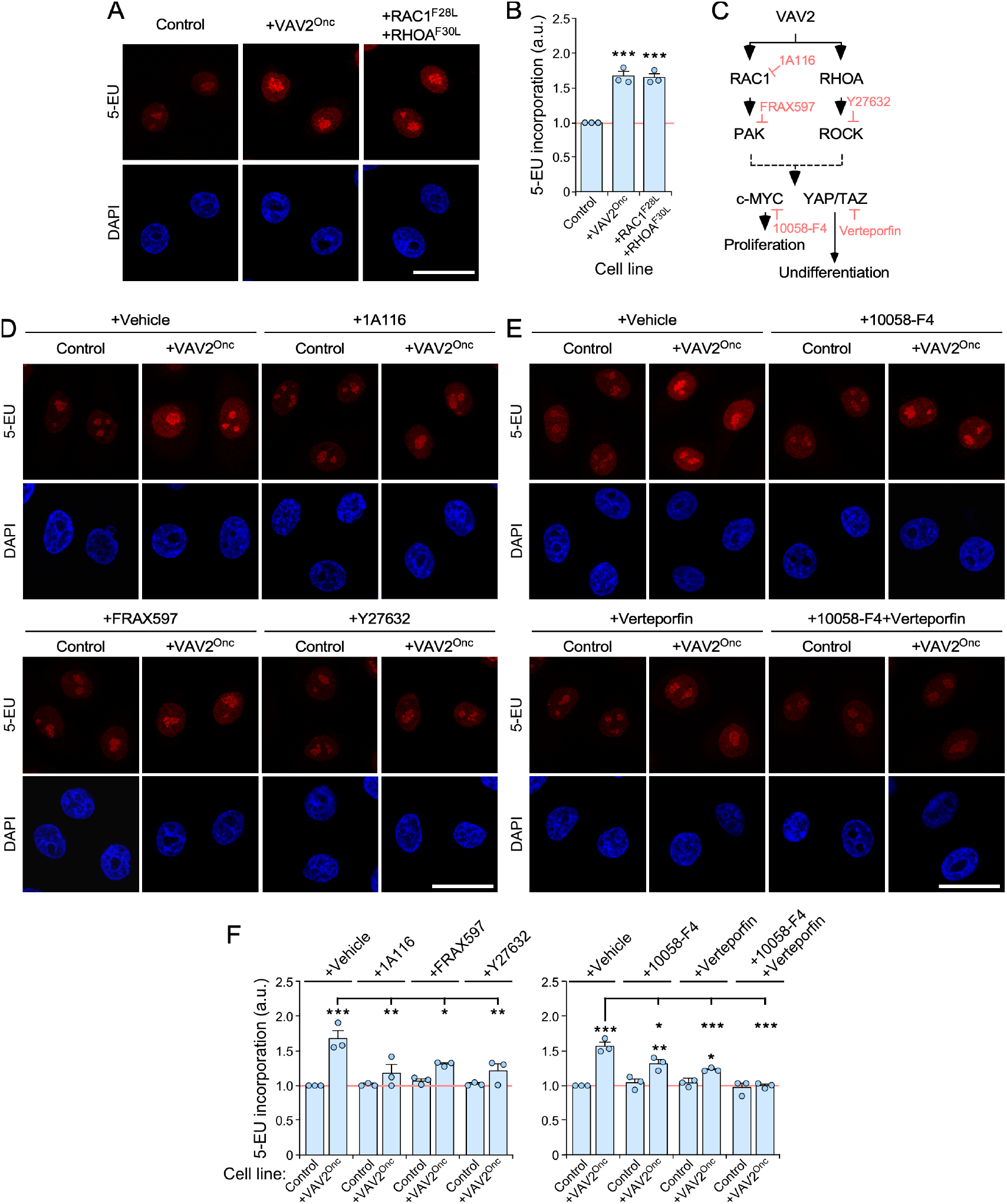
Mechanistic analysis of VAV2^Onc^-induced ribogenesis. (**A**) Representative images of 5-EU– (red color, top panels) and DAPI-labeled (blue color, bottom panel) human keratinocytes expressing the indicated proteins (top). Scale bar, 20 μm. **(B)** Quantification of the incorporation of 5-EU from the experiments shown in (A). ***, *P* < 0.0001 (ANOVA and Dunnett’s multiple comparison tests, *n* = 3 independent experiments). **(C)** Schematic representation of the VAV2^Onc^-mediated regulation of cell proliferation and undifferentiation in human keratinocytes based on previous work [1]. Inhibitors targeting specific VAV2 downstream elements are shown in light red. (**D** and **E**) Representative images of 5-EU– (red color) and DAPI-labeled (blue color) human keratinocytes expressing the indicated proteins and subjected to the culture conditions shown on the top. Scale bar, 20 μm. **(F)** Quantification of the incorporation of 5-EU from the experiments shown in (D) and (E). *, *P* < 0.05; **, *P* < 0.001; ***, *P* < 0.0001 vs controls or between the indicated experimental pairs (brackets) ANOVA plus Tukey’s HSD tests. *n* = 3 independent experiments. In (B) and (F), data represent the mean ± SEM. Source data for this figure are provided as a Source Data file.

We have previously shown using organotypic 3D cultures that the addition of the inhibitors for RAC1, PAK, ROCK and c-MYC blocks the epidermal hyperplasia induced by the stable expression of VAV2^Onc^ in keratinocytes (19) (see example in **Fig. 4A**). We found that this process is associated with a change in the distribution pattern of the 5.8S rRNA, as the inhibitor-treated 3D cultures of VAV2^Onc^-expressing keratinocytes show a control-like 5.8S rRNA immunoreactivity pattern (**Fig. 4A**, see quantitation in **Fig. 4B**, left). The same effect was observed in the organotypic cultures generated by VAV2^Onc^-expressing cells treated with c-MYC inhibitors (**Fig. 4C**, see quantitation in **Fig. 4B**, right). Unlike the rest of inhibitors, the addition of verteporfin promotes an extensive differentiation of keratinocytes located in the suprabasal layer (19). As a consequence, sections from these organotypic cultures keep that layer highly enlarged although, in this case, composed of differentiated cells (19) (**Fig. 4C**). The 5.8S rRNA immunoreactivity was totally absent from that differentiated layer (**Fig. 4C**; see quantitation in **Fig. 4B**, right). Same results were obtained in organotypic cultures of RAC1^F28L^+RHOA^F30L^-expressing cells treated with c-MYC or YAP/TAZ inhibitors (**Supplementary** Fig. 4; for quantitation, see **Fig. 4B**, right). Taken collectively, these results indicate that the upregulation of ribosome biogenesis is integrated in the same VAV2^Onc^-regulated signaling framework that promotes regenerative proliferation. They also indicate that ribogenesis is dually influenced by two independent mechanisms: (i) the c-MYC– and YAP/TAZ-dependent effect on RNA polymerase I activity, and (ii) the YAP/TAZ-mediated blockage of cell differentiation that favors a cell state associated with intrinsic high ribogenesis rates.

**FIGURE 4.**
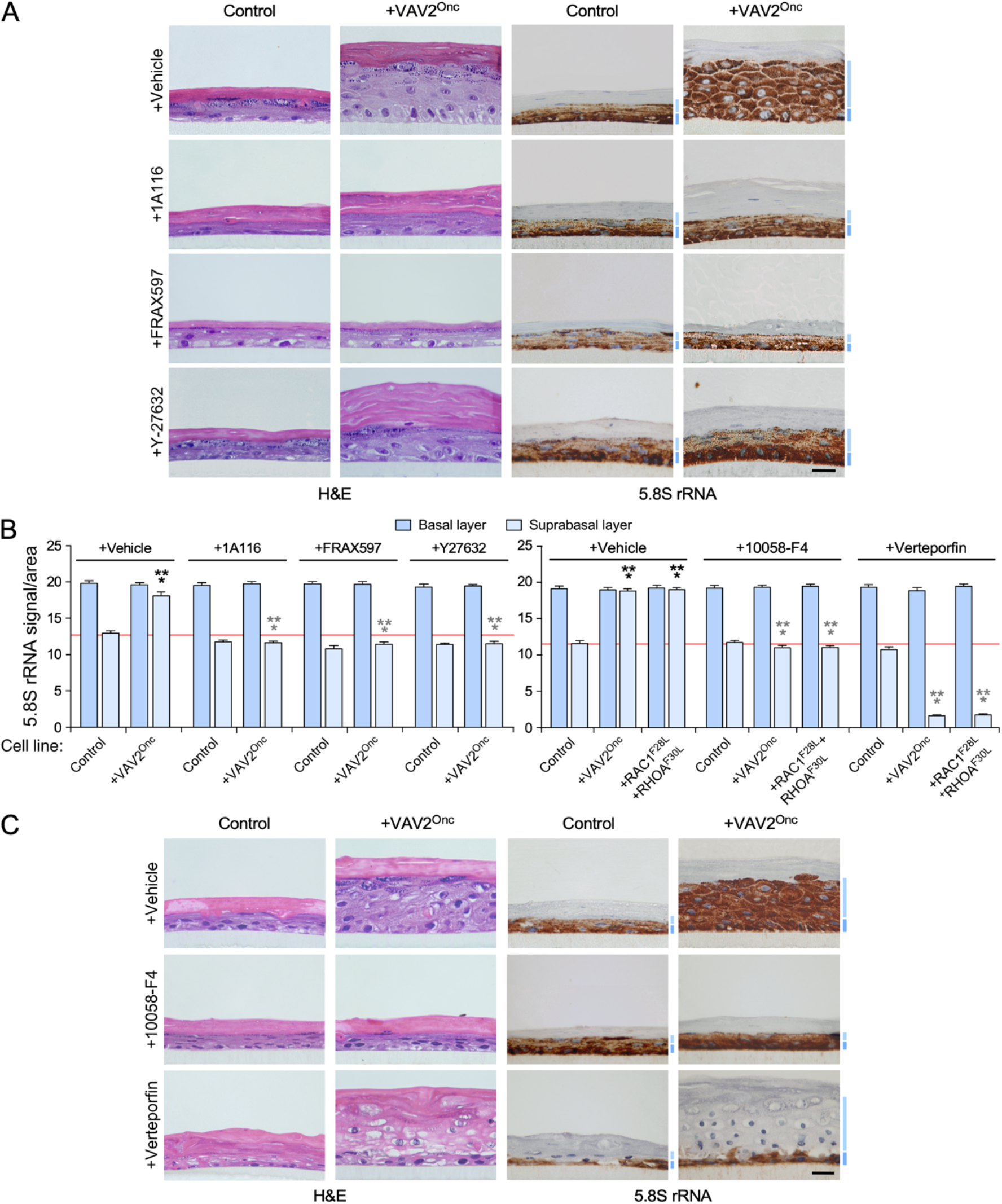
VAV2^Onc^-driven ribogenesis is PAK, ROCK, MYC, and YAP/TAZ-dependent. (**A** and **C**) Representative images of organotypic cultures of human keratinocytes expressing the indicated proteins (top) after staining with either hematoxylin-eosin (two left columns; A and C) or with an antibody to the 5.8S rRNA plus hematoxylin (two right panels; A and C). Dark and light blue bars represent the 5.8S rRNA immunoreactivity levels found in the basal and suprabasal epithelial layers, respectively. Scale bar, 10 μm. **(B)** Quantitation of the 5.8S rRNA immunoreactivity obtained in panels (A) and (C) of this figure as well as in the experiments shown in **Supplementary** Figure 6 (right panel). Black and gray asterisks indicate the *P* value of the indicated experimental values in untreated and treated cells when compared to the appropriate controls. ***, *P* < 0.0001 (ANOVA and Tukey’s HSD tests, *n* = 3 independent experiments). Data represent the mean ± SEM. Source data for this figure are provided as a Source Data file.

### Ribogenesis contributes to VAV2^Onc^-driven epidermal hyperplasia

Given that the concentrations of inhibitors selected in the previous experiments did not have any negative effect on the ribosome ribogenesis of control cells, we hypothesized that keratinocytes with upregulated VAV2 signaling could be highly dependent on high ribosome biogenesis rates to promote hyperplasia in organotypic cultures. To test this idea, we investigated the effect of a chemical inhibitor of RNA polymerase I (CX-5461) on the organotypic cultures generated by VAV2^Onc^ or c-MYC-expressing keratinocytes. Given the lethality associated with the total blockage of this polymerase, we selected a concentration of CX-5461 that did not impair the growth of both control and VAV2^Onc+E200A^-expressing cells (**Fig. 5A** to **C**). CX-4561 eliminated the hyperplasia (**Fig. 5A** and **B**) and restored a control cell-like distribution of the 5.8S rRNA in the epithelia formed by VAV2^Onc^-expressing keratinocytes (**Fig. 5A** and **C**). This result indicates that the tissue hyperplasia generated by these cells is highly dependent on high ribogenesis rates. In contrast, CX-4561 was much less effective when tested in 3D cultures of c-MYC-expressing cells (**Fig. 5A** and **B**). This is probably due to the higher rates of ribosome biogenesis in those cells, as inferred from the high levels of 5.8S rRNA immunoreactivity that are still detected in the CX-5461-treated epithelial structures formed by them (**Fig. 5A** and **C**). This idea is also consistent with the higher levels of 5-EU incorporation exhibited by these cells when compared to VAV2^Onc^-expressing keratinocytes (see above, **Fig. 2B** and **C**).

**FIGURE 5.**
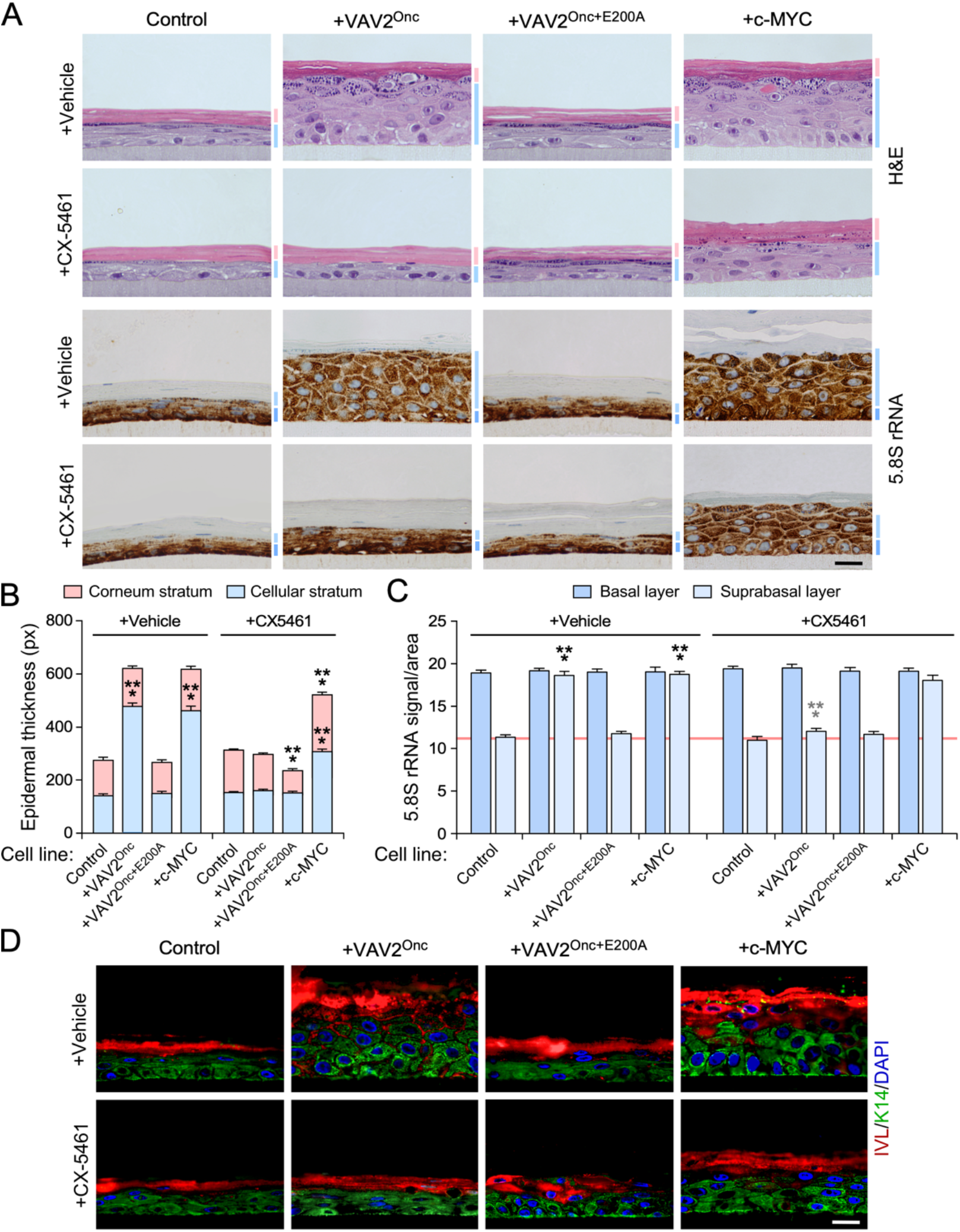
Ribogenesis is important for VAV2^Onc^-driven epidermal hyperplasia. (**A**) Representative images of organotypic cultures of human keratinocytes expressing the indicated proteins (top) and subjected to the culture conditions shown on the left that were stained with either hematoxylin-eosin (two top rows of panels) or with an antibody to the 5.8S rRNA plus hematoxylin (two bottom rows of panels). Dark and light blue bars represent the 5.8S rRNA immunoreactivity levels found in the basal and suprabasal epithelial layers, respectively. Scale bar, 10 μm. **(B** and **C)** Quantification of the thickness (B) and 5.8S rRNA immunoreactivity (C) of indicated cell layers obtained in the experiments shown in (A). In (C), Black and gray asterisks indicate the *P* value of the indicated experimental values when compared to control and vehicle-treated cells, respectively. ***, *P* < 0.0001 (ANOVA and Dunnett’s multiple comparison tests, *n* = 3 independent cultures). Data represent the mean ± SEM. Source data for this figure are provided as a Source Data file. px, pixel. **(D)** Representative images showing the expression of involucrin (IVL, red color) and keratin 14 (K14, green color) in organotypic cultures from indicated human keratinocytes (top) and culture conditions (left). Nuclei were counterstaining with DAPI (blue color). Scale bar, 10 μm. Similar results were obtained in two additional independent experiments (not shown).

The effect of CX-5461 is probably connected with an impact on the proliferation rather than the differentiation of VAV2^Onc^-expressing cells given that, unlike the case of the treatments with verteporfin (see above, **Fig. 4C**) (19), this inhibitor does not induce any histological sign of differentiation in the organotypic cultures (**Fig. 5A**). In line with this, we also observed that the distribution of involucrin-positive (a marker for differentiated cells) and keratin 14-positive (a marker for undifferentiated cells) is indistinguishable in the sections obtained from organotypic cultures generated by control cells, VAV2^Onc+E200A^-expressing cells, and CX-5461-treated VAV2^Onc^-expressing cells (**Fig. 5D**).

### The endogenous VAV2 pathway influences ribogenesis in oSCC cells

Having established that VAV2 signaling positively influences ribosome biogenesis rates in normal keratinocytes, we next investigated whether the activity of endogenous VAV2 could be also important for maintaining the ribogenic activity of already transformed oSCC cells. To this end, we utilized two previously described PDCs (VdH01, VdH15) from independent HPV^−^ oSCC cases that were stably transduced with control and *VAV2* short hairpin RNA (shRNA)-encoding lentiviral particles (19). In addition, we included control and *VAV2* knockdown derivatives of an oSCC cell line (SSC-25) that were generated following a similar strategy (19). We previously demonstrated using those experimental models that the endogenous WT VAV2 protein was important to maintain both the high proliferation and undifferentiated features of these three cell lines (19) (see examples in **Fig. 6A**). Upon staining of sections of the tissue structures formed by those cells in 3D cultures with antibodies to the 5.8S rRNA, we found that VdH01 and VdH15 cells are also highly dependent on endogenous VAV2 for maintaining high rates of ribosome biogenesis (**Fig. 6A** and **B**). The SSC-25 cell line, although highly dependent on VAV2 for overall growth (**Fig. 6A**) (19), maintain similar levels of the 5.8S rRNA in the absence and presence of endogenous VAV2 (**Fig. 6A** and **B**). The pathway that contributes to ribogenesis in oSCC PDCs is conserved with the one previously found in normal keratinocytes (see above, **Figs. 3** to **5**), as we observed that the inhibitors for RAC1, c-MYC or YAP/TAZ also promote a reduction in 5.8S rRNA immunoreactivity similar to that found in *VAV2* knockdown cells (**Fig. 6C** and **D**). These results indicate that the PDCs interrogated in this study are highly dependent on the activity of endogenous VAV2 signaling to maintain optimal rates of ribosome biogenesis. Moreover, the data obtained with SSC-25 cells suggest that, in some cases, the proliferative and ribogenic activity of cancer cells can be regulated by VAV2-dependent and independent pathways, respectively.

**FIGURE 6.**
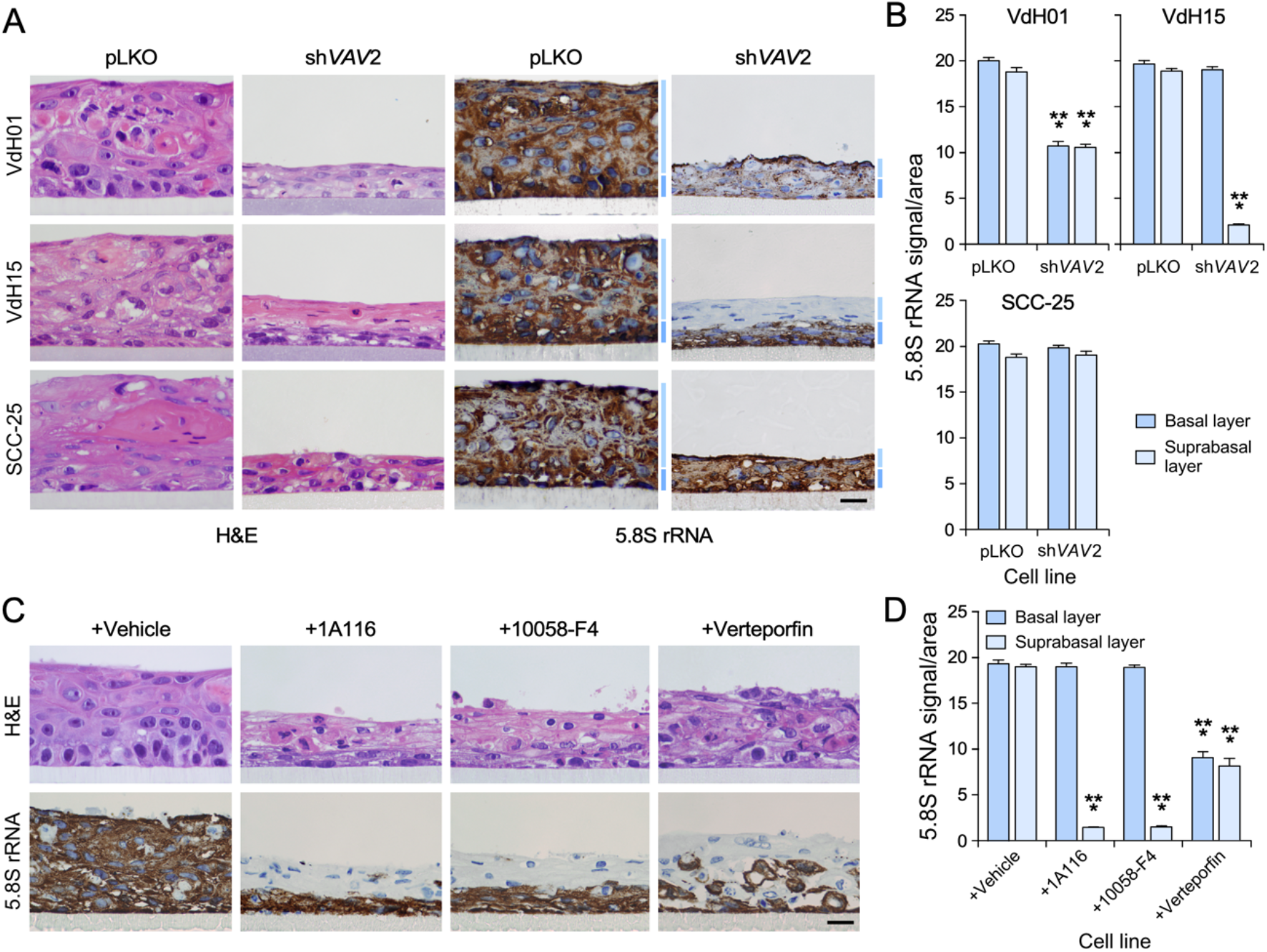
The endogenous VAV2 pathway influences ribogenesis in oSCC cells. (**A**) Representative images of organotypic cultures of indicated control and *VAV2*-knockdown oSCC cells that stained with either hematoxylin-eosin (two left columns) or with an antibody to the 5.8S rRNA plus hematoxylin (two right panels). Dark and light blue bars (right) indicate cell layers analogous to the basal and suprabasal strata formed by normal keratinocytes, respectively. Scale bar, 10 μm. pLKO, control cells containing an empty lentiviral vector. **(B)** Quantitation of the 5.8S rRNA immunoreactivity obtained in panel (A). ***, *P* < 0.0001 (Student’s t-test, *n* = 3 independent experiments). **(C)** Representative images of organotypic cultures of VdH15 cells treated as indicated (top) that were stained with either hematoxylin-eosin (top panels) or with an antibody to the 5.8S rRNA plus hematoxylin (bottom panels). Scale bar, 10 μm. **(D)** Quantitation of the 5.8S rRNA immunoreactivity obtained in panel (C). ***, *P* < 0.0001 (Student’s t-test, *n* = 3 independent experiments). In (B) and (D), data represent the mean ± SEM. Source data for this figure are provided as a Source Data file.

### Ribosome ribogenesis is a therapeutic Achilles’ heel for VAV2-dependent oSCCs

Previous reports have shown that the elimination of specific ribosome biogenesis factors (HEATR1, NOB1, PES1, RIOK2) impairs the proliferation and malignant traits of oSCC cell lines (22–25). This is not entirely surprising given that the elimination of these proteins is lethal (13). Likewise, it has been shown that the inhibition of RNA polymerase I *per se* or in combination with mTORC reduces the *in vivo* tumorigenicity of an oSCC cell line using xenotransplants experiments (11). To further assess this issue, we investigated the effect of CX-5461 on VdH01, VdH15, and SSC-25 cells using organotypic 3D cultures. As in previous experiments, we selected a concentration of the inhibitor that did not impair the growth of normal keratinocytes to avoid non-specific effects derived from the total shutdown of this essential process. CX-5461 reduced the growth of the two oSCC PDCs used in our study (**Fig. 7A** and **B**), a process that was associated, as expected, with the reduction in 5.8S rRNA immunoreactivity levels in both cases (**Fig. 7A** and **C**). By contrast, the RNA polymerase I inhibitor did not elicit any statistically significant effect on the 3D growth of SSC-25 cells (**Fig. 7A** and **B**). No changes in 5.8S rRNA immunoreactivity were observed in this case either (**Fig. 7A** and **C**), suggesting that these cells might have higher RNA polymerase I activity than PDCs. We did not find any overt signs of differentiation in the sections obtained from CX-5461-treated cultures (**Fig. 7D**).

**FIGURE 7.**
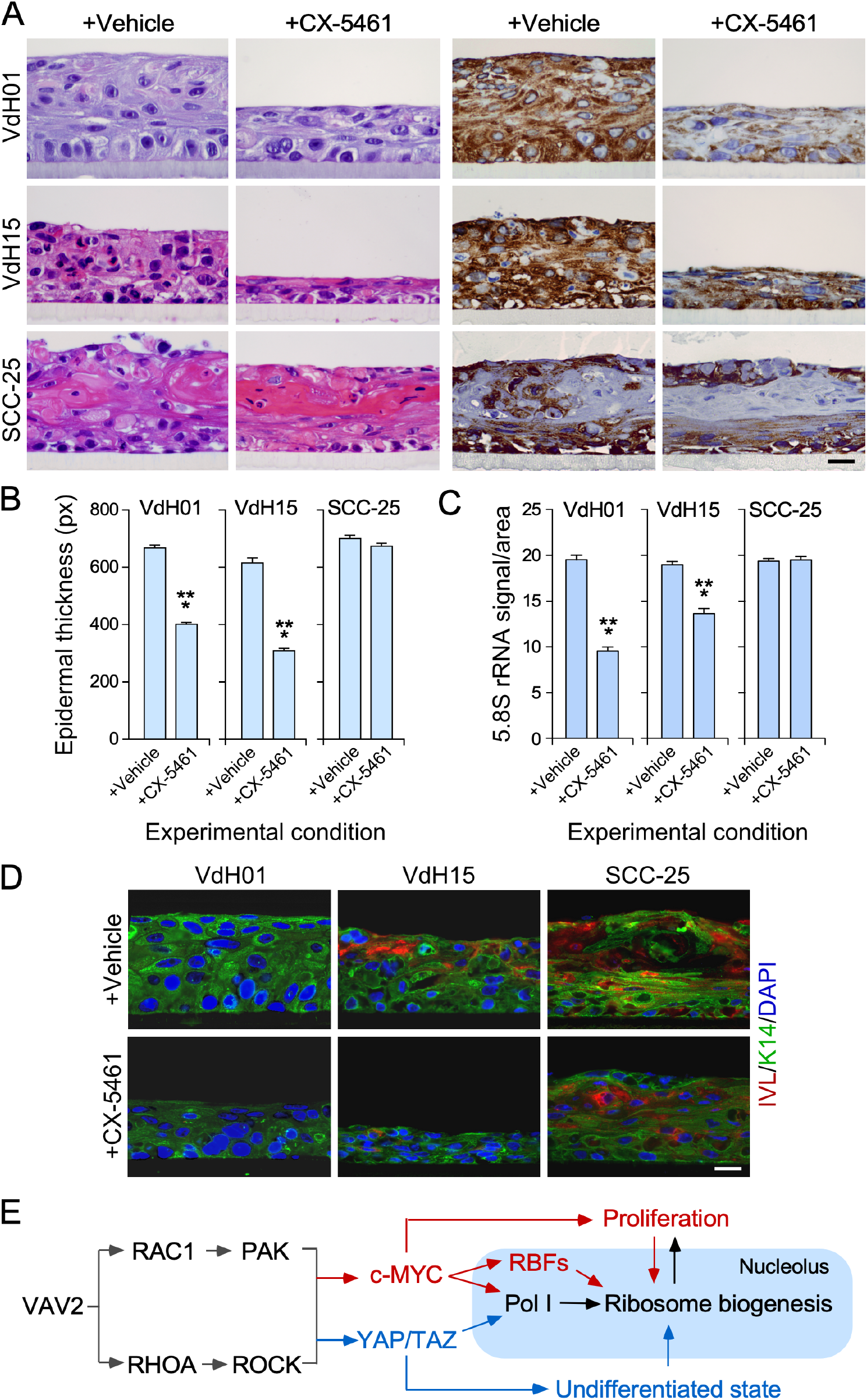
oSCC cells are sensitive to high rates of ribosome ribogenesis. (**A**) Representative images of organotypic cultures of indicated oSCC cells (left) that were treated with either vehicle solution or CX-5461 (top). Sections were stained with hematoxylin-eosin (two left columns) or with an antibody to the 5.8S rRNA plus hematoxylin (two right columns). Scale bar, 10 μm. **(B** and **C)** Quantitation of the thickness of the epithelium (B) and the 5.8S rRNA immunoreactivity (C) obtained in experiments shown in (A). ***, *P* < 0.001 (Student’s *t*-test, *n* = 3 independent experiments). Data represent the mean ± SEM. Source data for this figure are provided as a Source Data file. **(D)** Representative images of organotypic models from indicated oSCC cell lines (top) and culture conditions (left) that were stained with antibodies to either involucrin (IVL, red color) or keratin 14 (K14, green color) and counterstained with DAPI (blue color). Scale bar, 10 μm. Similar results were obtained in two additional experiments (not shown). **(E)** Proposed model for the regulation of ribosome biogenesis by the VAV2 catalysis-dependent pathways in normal and transformed keratinocytes. Pol I, RNA polymerase I; RBFs, ribosome biogenesis factors. The nucleolus is shaded in blue. The bidirectional arrows between proliferation and ribosome biogenesis indicate that both processes influence each other. The unidirectional arrow between the undifferentiated state and ribosome biogenesis indicates that the former affects the latter but not *vice versa*.

## DISCUSSION

There is still very scant information on possible signaling connections between RHO GTPases and ribogenesis in both normal and cancer cells. In this work, we have demonstrated that the RHO GEF VAV2 plays important roles in the upregulation and maintenance of ribosome biogenesis levels in normal keratinocytes and oSCC PDCs, respectively. This result indicates that the correlation seen in bioinformatics analyses between the expression levels of the *VAV2* mRNA and the ribosome biogenesis-related gene signatures in patient samples probably reflects a direct functional connection between those two routes rather than just a mere statistical correlation.

Mechanistic analysis performed using both 2D and 3D cultures indicate that, unlike the case of ECT2 and ARHGAP30 (17, 18, 26), the connection established between VAV2 and ribogenesis involves a more canonical signaling pathway that entails the VAV2-mediated stimulation of the GTPases RAC1 and RHOA, the proximal effectors PAK and ROCK, and the transcriptional factors c-MYC and YAP/TAZ. The stimulation of both c-MYC and YAP/TAZ leads to increased production of the 47S pre-rRNA precursor in the nucleolus through the stimulation of RNA polymerase I activity (**Fig. 7E**). In addition, we believe that the YAP/TAZ complex also contributes indirectly to this process via maintaining a highly cell undifferentiated state that is compatible with full ribogenic activity (**Fig. 7E**). It is likely that these two inputs are further boosted by the large collection of ribosome biogenesis factors that become upregulated during VAV2^Onc^-driven epidermal hyperplasia. Although we have not mechanistically dissected this part of the equation, previous studies suggest that this could be mediated by c-MYC (12) (**Fig. 7E**). Given the level of conservation of this route in primary keratinocytes and oSCC PDCs, it is likely that the mechanism reported here for the VAV2-mediated regulation of ribosomal will be also operative in other subtypes of hnSCC.

Interestingly, the regulation of ribosome biogenesis in basal cells seem to be different from the mechanistic model shown here for the hyperplasic keratinocytes (**Fig. 7E**). This is based on several observations: (i) the 5.8S rRNA immunoreactivity is mostly concentrated in the basal layer of the epithelial structures formed by control cells; (ii) this immunoreactivity is preserved in organoid cultures from VAV2^Onc^-expressing or RAC1^F28L^+RHOA^F30L^-expressing keratinocytes that have been treated with inhibitors for downstream VAV2 signaling elements; and (iii) it is also maintained when 3D cultures from both control and VAV2^Onc^– or RAC1^F28L^+RHOA^F30L^-expressing keratinocytes are treated with CX-5461. This latter result suggests that basal cells in all those cultures have higher levels of RNA polymerase I activity than the suprabasal counterparts. The VAV2–independent mechanism that controls ribogenesis in basal cells remains to be determined. However, it is likely that it is ECT2– and RAC1-independent given that the RAC1 inhibitor used in our study did not affect the 5.8S rRNA immunoreactivity levels exhibited by those cells.

VAV2 signaling plays critical roles in the regenerative proliferation of keratinocytes, a feature that is associated with poor patient prognosis (19). We surmise therefore that the targeting of signaling elements of this pathway can be potentially interesting to treat this type of tumors. The results presented here further emphasize this idea, as they have expanded the influence of this signaling route to the regulation of ribosome biogenesis. This new VAV2-dependent response certainly contributes to the malignancy of oSCC cancer cells, as demonstrated by the negative effects of the CX-5461 inhibitor on the growth of PDCs in 3D cultures. We have seen no interference with the same parameters in control cells at the concentration of inhibitor used, thus suggesting that there are therapeutic windows in which the activity of healthy cells could be preserved. Interestingly, we have observed that the dependency on ribosome biogenesis can vary depending on the oSCC cell used. For example, we have found that ribogenesis is both VAV2– and CX-5461-independent in the case of the SCC-25 cell line. The growth of SCC-25 cells is VAV2-dependent (19), suggesting that they have acquired specific signaling and/or genetic alterations that have uncoupled the regulation of ribogenesis from the rest of VAV2-dependent processes.

How to target this new VAV2-regulated process in oSCC and other hnSCC subtypes? One possibility is to target VAV2 itself via either standard or PROTAC-based inhibitory approaches. This avenue has been a problematic so far, given the unfruitful avenues made before with many RHO GEFs (27). Another strategy is to focus on more druggable downstream elements such as c-MYC or the YAP/TAZ complex (28–30) (e.g., clinical trials NCT05100251, NCT05228015, NCT04665206). This option has the advantage that it can kill regenerative proliferation and ribogenesis at the same time in tumors. Another plausible option is targeting RNA polymerase I itself (**Fig. 7E**), an avenue that has already been demonstrated in mouse models (11, 31, 32) and is being tested in ongoing clinical trials (NCT04890613). Further studies will be needed to pinpoint the best therapeutic strategies to block this new pathway in hnSCC and, perhaps, in other VAV2-dependent SCC subtypes.

## EXPERIMENTAL PROCEDURES

### Ethics statement

All animal work was performed in accordance with protocols approved by the Bioethics committee of Salamanca University and the animal experimentation authorities of the autonomous government of Castilla y León (Spain). The use of PDCs was conducted according to methods and *a priori* informed patient consent policies approved by the Bioethics committees of the Vall d’Hebron Research Institute.

### Plasmids

The plasmid encoding EGFP-VAV2^Onc^ (pNM115) was described before (19). For its construction, the cDNA encoding VAV2^Onc^ (Σ1-186) was liberated from plasmid pKES19 (33) by digestion with BstXI, filled-in, and cloned into the SmaI-linearized pEGFP-C2 vector (Clontech, Catalog No. 632481). Luciferase activity was conducted using a plasmid encoding luciferase under the regulation of the rRNA promoter (provided by L.-L. Chen, Institute of Biochemistry and Cell Biology, Chinese Academy of Sciences, University of Chinese Academy of Sciences, 200031 Shanghai, China) and pRL-SV40 (*Renilla* luciferase, obtained from Promega, Catalog No. E2231). All plasmids were DNA sequence verified.

### Cells

Primary human keratinocytes (Ker-CT cell line, immortalized by the ectopic expression of both TERT and CDK4) were obtained from the American Type Culture Collection (Catalog No. CRL-4048). These cells were cultured in CnT-Prime medium (CELLnTEC, Catalog No. CnT-PR) and transfected in KGM-Gold medium (Lonza, Catalog No. 00192060). oSCC PDCs (VdH01, VdH15) were generously provided by S.A. Benitah (Institut Reserca Biomédica, Barcelona, Spain) and described elsewhere (19, 34). VdH01 cells were cultured in FAD^+^ medium, which is a combination of 75% DMEM (Gibco, Catalog No. 21969) and 25% Ham’s F12 medium (Thermo Fisher, Catalog No.11765054) that were supplemented with 10% fetal bovine calf serum (Gibco, Catalog No. 10270106), 2 mM L-glutamine (Gibco, Catalog No. 25030024), 1.8 x 10^−4^ M adenine (Sigma-Aldrich, Catalog No. A2786–5G), 0.5 μg/mL hydrocortisone (Sigma-Aldrich, Catalog No. H4001-1G), 5 μg/mL insulin (Thermo Fisher, Catalog No. 12585014), 10 ng/mL epidermal growth factor (PreproTech, Catalog No. AF-100-15) and 10^−10^ M cholera toxin (Sigma-Aldrich, Catalog No. C8052-5MG). VdH15 cells were grown in KSFM medium supplemented with 25 μg/mL BPE and 0.5 ng/ml epidermal growth factor. 3D cultures were carried out as indicated above. SCC-25 cells were provided by S.A. Benitah and cultured in KSFM medium supplemented with 25 μg/mL BPE and 0.5 ng/mL epidermal growth factor. Derivatives from all those cells expressing or lacking the indicated proteins were generated and validated in a previous study from our lab (19). When appropriate, inhibitors for indicated signaling elements were used. Those included: 1A116 (500 nM) (35, 36), FRAX597 (5 nM, Selleckchem, Catalog No. S7271), Y-27632 (1 μM, Selleckchem, Catalog No. S1049), 10058-F4 (500 nM, Selleckchem, Catalog No. S7153), verteporfin (100 nM, Selleckchem, Catalog No. S1786), and CX-5461 (100 nM, Selleckchem, Catalog No. S2684). Concentration of inhibitors were chosen based on their minimal effect on control cells in 2D and 3D culture conditions.

### Mouse models

*Vav2*^Onc/Onc^ knock-in mice and appropriate controls have been described elsewhere (37). Animals were kept in ventilated rooms in pathogen-free facility of the University of Salamanca under controlled temperature (23°C), humidity (50%), and illumination (12-hour-light/12-hour-dark cycle) conditions.

### *In silico* analyses of mouse expression microarray data

The functional annotation of the VAV2^Onc^-dependent transcriptome was reported before using the Gene Expression Omnibus (GEO) dataset GSE124019 [https://www.ncbi.nlm.nih.gov/geo/query/acc.cgi?acc=GSE124019] (19). Gene set enrichment analyses (https://www.gsea-msigdb.org/gsea/index.jsp) were performed using the same GEO dataset using gene set permutations (*n* = 1000) for the assessment of significance and signal-to-noise metric for ranking genes. Protein interaction networks were built using the Cytoscape software (https://cytoscape.org, National Resource for Network Biology). To evaluate the expression of the VAV2^Onc^ gene signature for ribosome biogenesis factors across normal, dysplastic and tumoral samples, the enrichment score was calculated using ssGSEAs (https://www.genepattern.org/modules/docs/ssGSEAProjection/4#gsc.tab=0). To this end, we used the GEO GSE30784 dataset (*n* = 229 samples) [https://www.ncbi.nlm.nih.gov/geo/query/acc.cgi?acc=GSE30784] (38). This dataset lacks information on HPV status, although the percentage of HPV^−^ cases contained in this dataset has been estimated to be in the 75% range (38).

Overall survival analyses were performed through Kaplan-Meier estimates according to the expression level of indicated signatures using the GEO GSE41613 [https://www.ncbi.nlm.nih.gov/geo/query/acc.cgi?acc=GSE41213] dataset. This dataset was selected because it contained information on long-term survival, HPV status, and other clinical criteria of patients. It also contain a number of samples (*n* = 97 cases, all of them HPV^−^) that were compatible with proper statistical analyses (39). The median of the expression distribution of the indicated gene signature was used to establish the low and high expression groups and, subsequently, the Mantel-Cox test was applied to statistically corroborate the differences seen between the two survival distributions. The survival scores for the *EGFR* mRNA, the *VAV2* mRNA, and other VAV2^Onc^-regulated gene signatures were calculated in a previous study using the same method and GEO dataset (19).

### Isolation of primary mouse keratinocytes

Primary keratinocytes were isolated as previously described (19). Briefly, the skin from euthanized neonatal mice of indicated genotypes was incubated with 250 units/mL dispase (Roche, Catalog No. 04942078001) in KSFM medium (Thermo Fisher, Catalog No. 17005-042) for 16 h at 4 °C to separate the epidermis from the dermis. The epidermis was then treated with accutase (CELLnTEC, Catalog No. CnT-Accutase-100) for 30 min at 37°C to release the keratinocytes. The isolated cells were then cultured in KSFM medium supplemented with 20 nM CaCl_2_, 25 μg/mL BPE and 0.25 ng/mL EGF (Thermo Fisher, Catalog No. 37000-015).

### Three-dimensional organotypic cultures

Mouse (5 x 10^5^ cells) and human (2 x 10^5^ cells) keratinocytes were seeded onto 12 mm diameter inserts (Millipore, Catalog No. PIHP01250) and cultured in CnT-Prime medium. After two days, the medium was replaced with CnT-PR 3D-Barrier (CellnTec, Catalog No. CnT-PR-3D) and 16 h later the airlift was performed according to manufacturer’s instructions. 3D cultures were maintained in CnT-PR 3D-Barrier for 12 days and ultimately fixed in 4% paraformaldehyde to be processed for immunohistochemical analysis. During the final seven days of the culture, cells were treated with the appropriate vehicles and inhibitors, including 1A116 (500 nM), FRAX597 (5 nM), Y-27632 (1 μM), 10058-F4 (500 nM), verteporfin (100 nM), and CX-5461 (100 nM). Concentration of inhibitors were chosen based on their minimal effect on control cells.

### Histological and immunohistochemical studies

Tissue sections were either stained with either hematoxylin-eosin or exposed to Tris EDTA [pH 8.0] for heat-induced antigen unmasking and subsequent incubation for 40 min with the appropriate primary antibody to 5.8S rRNA (1:1500 dilution, Santa Cruz, Catalog No. sc-33678), involucrin (1:100 dilution, Sigma-Aldrich, Catalog No. I9018) or keratin 14 (1:300 dilution, Biolegend, Catalog No. 905301). Immunohistochemical staining was carried out using a Ventana Discovery Ultra instrument (Roche, Catalog No. 05987750001). For standard staining, the Discovery OmniMap anti-rabbit horse radish peroxidase detection system (Roche, Catalog No. 760-4311) was used for detection as specified by the manufacturer. For immunofluorescent studies, sections were incubated for 1 h with appropriate secondary antibodies to either rabbit or mouse IgGs labeled with Alexa Fluor 488 (1:200 dilution, ThermoFisher, Catalog No. A21206) and Cy3 (1:200 dilution, Jackson ImmunoResearch, Catalog No. 115-165-146). For cell nuclei staining, sections were incubated with 4′,6-diamidino-2-phenylindole dihydrochloride (Sigma-Aldrich, Catalog No. D9542) for 5 min. Immunohistochemical signals were quantified using the Fiji software.

### Determination of nucleolar parameters

Exponentially growing cells were seeded onto 10 mm glass coverslips previously treated with poly-L-lysine (Sigma-Aldrich, Catalog No. F8775). After 48 h, cells were fixed with 4% paraformaldehyde in phosphate-buffered saline solution for 15 min, and permeabilized with 0.25 % Triton (Sigma-Aldrich, Catalog No. X100) in TBS-T [25 mM Tris-HCl (pH 8.0), 150 mM NaCl, 0.1% Tween-20 (Sigma-Aldrich, Catalog No. P7949)] for 10 min. Coverslips were then blocked with 2% bovine serum albumin in TBS-T for 30 min and incubated with a primary antibody to nucleophosmin (1:50 dilution, Invitrogen, Catalog No. 32-5200) in a moist chamber for 2 h. Next, cells were incubated with the corresponding secondary antibody (1:500, Invitrogen, Catalog No. A28175) for 30 min and stained with 4′,6-diamidino-2-phenylindole dihydrochloride for 5 min to visualize the nuclei. The coverslips were mounted onto glass slides using Mowiol medium and images were captured using a Leica TCS-SP8 microscope. The area of the nucleoli was measured using the Fiji software.

### 5-ethynyl uridine incorporation assays

Newly synthesized rRNA was examined using the Click-iT RNA Alexa Fluor 594 Imaging Kit (ThermoFisher, Catalog No. C10330). To this end, keratinocytes were plated onto poly-L-lysine-coated coverslips, cultured for 48 h, and treated with 1 mM 5-ethynyl uridine (ThermoFisher, Catalog No. C10330) for 20 min. Cells were then fixed with 4% paraformaldehyde in phosphate-buffered saline solution for 15 min, permeabilized with 0.25 % Triton X-100 in TBS-T for 10 min and blocked with 2% bovine serum albumin in TBS-T for 30 min. Nascent rRNA was detected using Alexa Fluor 594 according to the manufacturer’s protocol. Upon labeling, nucleoli were stained with 4′,6-diamidino-2-phenylindole dihydrochloride for 5 min and coverslips were mounted on slides. When indicated, cells were treated with the corresponding vehicles and inhibitors for 24 h. Inhibitors used included FRAX597 (5 nM), Y-27632 2HCl (1 µM), 1A116 (500 nM), 10058-F4 (500 nM), and verteporfin (200 nM). Concentrations of inhibitors were selected based on the induction of minor effect in the organotypic structures formed by control cells. Cell images were acquired using Leica TCS-SP8 microscope and the signal intensity was measured using the Fiji software.

### Luciferase assay for detection of DNA polymerase I activity

Exponentially growing cells were transiently transfected using Fu-GENE HD reagent (Promega, Catalog. No E2311) with: (i) 80 ng of the pRL-SV40 vector encoding the Renilla luciferase gene used as an internal control for transfection efficiency; (ii) 2 μg of the reporter plasmid containing the firefly luciferase gene under the regulation of the rRNA promoter; and (iii) 2 μg of the indicated EGFP-derived plasmids. After 36 h, cells were lysed with Passive Lysis Buffer (Promega, Catalog. No. E1960) and luciferase activity determined using the Dual Luciferase Assay System (Promega, Catalog No. E1960). In all cases, the ratio of the firefly luciferase/renilla luciferase activity was calculated and normalized to the control.

### Western blot analyses

Exponentially growing cells were washed with chilled phosphate buffered saline solution and then directly lysed in RIPA buffer at 4 °C. Extracts were precleared by centrifugation at 13 200 rpm for 10 min at 4°C, denatured by boiling in SDS-PAGE sample buffer, separated electrophoretically, and transferred onto nitrocellulose filters using the iBlot Dry Blotting System. Membranes were blocked as above and then incubated overnight at 4 °C with appropriate antibodies to GFP (1:1000 dilution; Clontech, Catalog No. 1. 632381) and tubulin α (1:2000 dilution; Calbiochem, Catalog No. CP06). After three washes with TBS-T to eliminate the primary antibody and immunoreacting bands revealed using a standard chemiluminescent method (Thermo Fisher Scientific, Catalog No. 32106).

### Northern blot analyses

Total RNAs were extracted using the Trizol method (TRI reagent, Ambion, Catalog No. AM9738) and quantified using a NanoDrop Spectrophotometer. Norther blot analyses were carried out following standard procedures after separation of RNA samples in 1.2% agarose/formaldehyde gels (40). The sequences of the oligonucleotides used as probes included: 5’-CCT CGC CCT CCG GGC TCC GGG CTC CGT TAA TGA TC-3’ (forward, *5’-ITS1*), 5’-GAT CAT TAA CGG AGC CCG GAG CCC GGA GGG CGA GG-3’ (reverse, *5’-ITS1*), 5’-CTG CGA. GGG AAC CCC CAG CCG CGC A-3’ (forward, *ITS2*), 5’-TGC GCG GCT GGG GGT TCC CTC GCA G-3’ (reverse, *ITS2*). RNA levels were quantified using the Fiji software.

### Statistical analyses

Statistics were calculated using GraphPad Prism 8.0 (Dotmatics) The number of biological replicates (n), the type of statistical test applied, and the statistical significance for each experiment are indicated in the corresponding figure legend. Data normality was tested using the Shapiro-Wilk test. Parametric distributions were analyzed using Student’s *t*-test (when comparing two experimental groups) or one-way ANOVA followed by either Dunnett’s (when comparing more than two experimental groups with a single control group) or Tukey’s HSD test (when comparing more than two experimental groups with every other group). In all cases, values were considered significant when *P* ≤ 0.05. Data obtained are expressed as the mean ± SEM. Heatmaps were generated using the heatmap3 *R* package.

## MATERIALS AVAILABILITY

All relevant data are available from the corresponding author upon reasonable request. A Materials Transfer Agreement could be required in the case of potential commercial applications.

## DECLARATION OF INTEREST

The authors declare no competing interests.

## AUTHOR CONTRIBUTIONS

N.F.-P. participated in all experimental work, analyzed data, and contributed to artwork design and manuscript writing. L.F.L.-M. carried out *in silico* analyses, generated cell lines, and carried out organotypic cultures. J.M.G.-P. and J.P.R. carried out immunohistochemical analyses. M.D. and X.R.B. conceived the work, analyzed data, wrote the manuscript, and carried out the final editing of figures.

## ACKNOWLEDGEMENTS

We thank A. Abad and C. García-Macías for lab and pathology work, respectively. The X.R.B.’s project leading to these results has received funding from Worldwide Cancer Research (14–1248), the Castilla-León government (CSI145P20, CLC-2017-01), grants cofounded by MCIN/AEI/10.13039/501100011033/ plus the European Research Development Fund “A way of making Europe” of the European Union (PID2021-122666OB-I00), “la Caixa” Banking Foundation (HR20-00164), and the Programa Excelencia of the Fundación Científica AECC 2022 (EPAEC222641CICS). M.D.’s work has been supported by a grant cofounded by MCIN/AEI/10.13039/501100011033/ plus the European Research Development Fund “A way of making Europe” of the European Union (PID2020-118378GB-I00). The authors’ institution was supported by the Programas de Apoyo a Planes Estratégicos de Investigación de Estructuras de Investigación de Excelencia of the Castilla-León government (CLC-2017-01 and CL-EI-2021-02) that were both cofounded by the European Research Development Fund. N.P. contract has been mostly supported by funding from the Spanish Ministry of Universities (FPU17/03912) and, subsequently, by the HR20-00164 grant (see above). L.F.L.-M. contract has been mostly supported by funding from the Spanish Ministry of Education, Culture and Sports (FPU13/02923) and, subsequently, by the CLC-2017-01 grant.

## Notes

### Competing Interest Statement

The authors have declared no competing interest.

