## Supplemental Information for "VAV2-Dependent Regulation of Ribosome Biogenesis in Keratinocytes and Oral Squamous Cell Carcinoma"

by

Natalia Fernández-Parejo, L. Francisco Lorenzo-Martín *et al*.

This PDF file includes:

Supplementary Figures 1 to 5 and legends

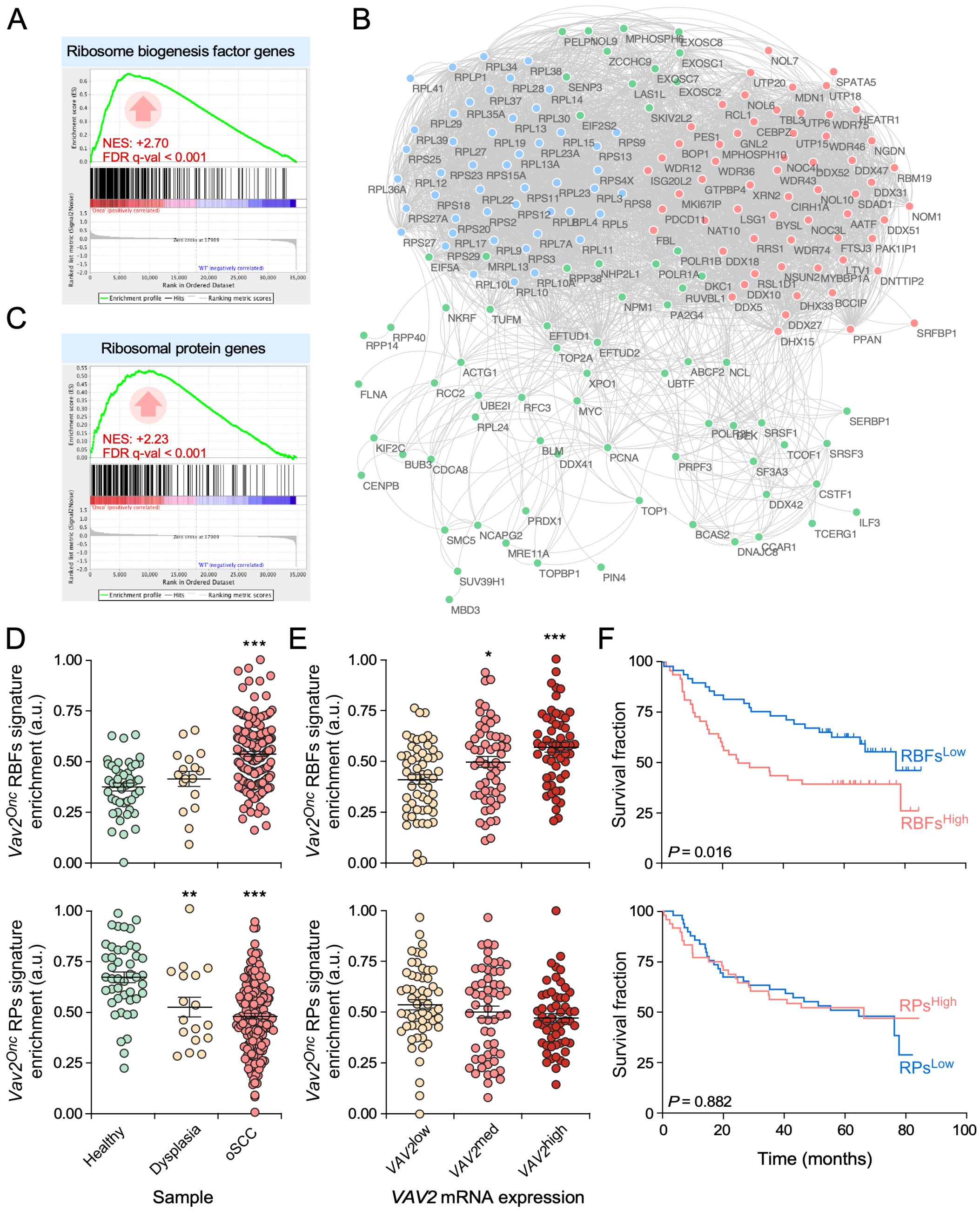


**SUPPLEMENTARY FIGURE 1. Ribogenesis correlates with both VAV2 signaling and hnSCC patient prognosis**

**(A and C)** GSEA enrichment plots showing the upregulation of gene sets for ribosome biogenesis factors (RBF, A) and ribosomal proteins (RP, B) in the VAV2$^{Onc}$-dependent transcriptome previously identified in the skin of mice. The normalized enrichment scores (NES) and false discovery rate q-values (FDR q-val) are indicated within each graph. Positive enrichments are indicated with upward arrows.

**(B)** Protein interaction network containing the 180 leading-edge genes obtained in (A) and (B). The red (right) and the blue (left) nodes constitute the VAV2$^{Onc}$ RBF and VAV2$^{Onc}$ RP signature, respectively. The green nodes represent other regulators of ribosome biogenesis.

**(D and E)** Dot plots showing the enrichment of the VAV2$^{Onc}$-regulated RBF (D and E, top panels) and VAV2$^{Onc}$-regulated RP (D and E, bottom panels) gene signatures according to patient sample (D) and *VAV2* mRNA expression levels within oSCC tumors (E). *, $P < 0.05$; **, $P < 0.01$; ***, $P < 0.001$ (ANOVA and Dunnett's multiple comparison tests). Data represent the mean ± SEM. Source data for this figure are provided as a Source Data file.

**(F)** Kaplan-Meier plots showing the survival rates of oSCC patients that were stratified according to the high and low expression levels of the VAV2$^{Onc}$-regulated RBF (top) and RP gene (bottom) signatures. The Mantel-Cox test *P* value is indicated for each transcript.

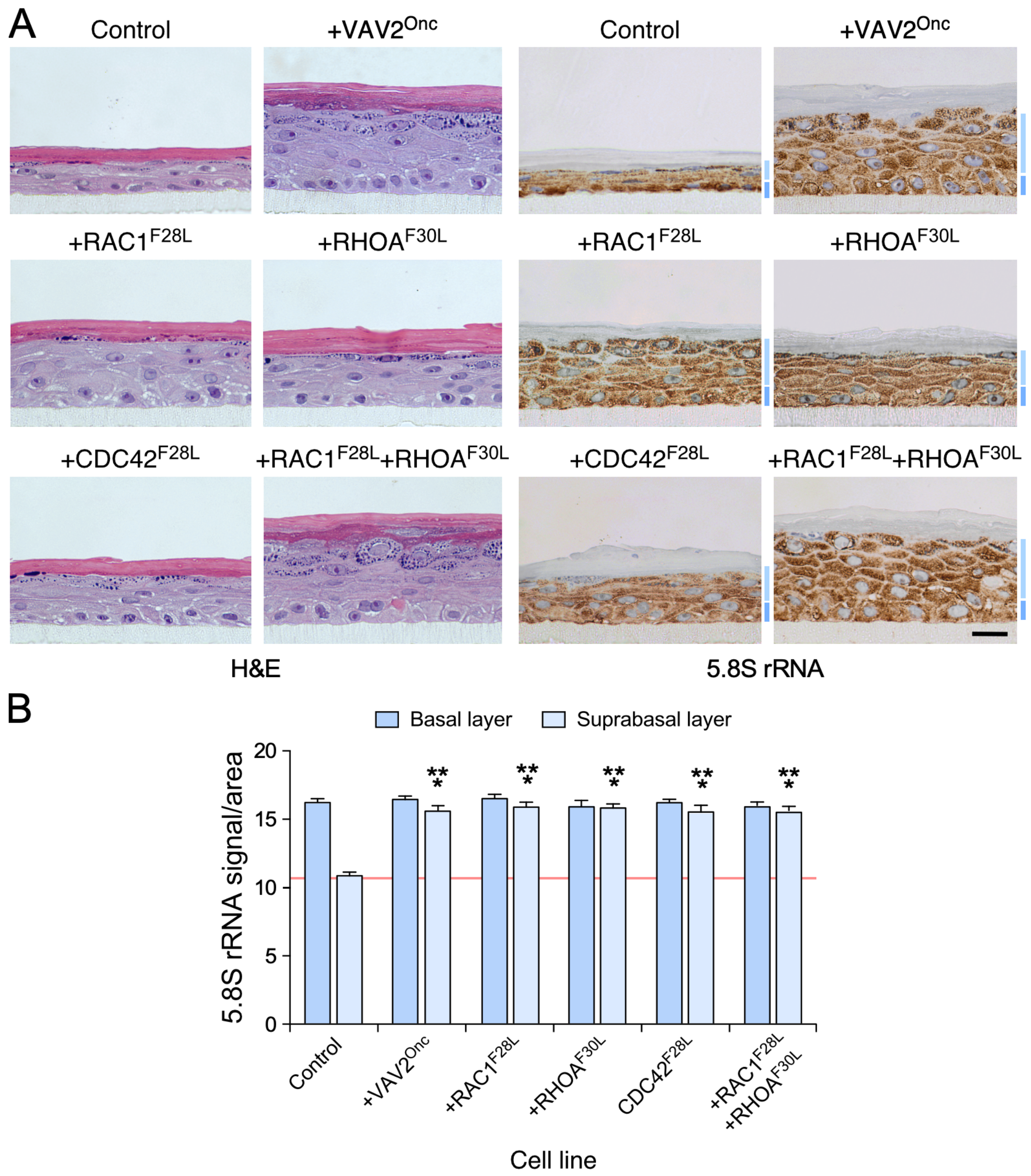


**SUPPLEMENTARY FIGURE 2. Activated RHO proteins also stimulated ribosome biogenesis**

**(A)** Representative images of organotypic cultures of human keratinocytes expressing the indicated proteins (top) that were stained with either hematoxylin-eosin (H&E, two left columns) or with an antibody to the 5.8S rRNA plus hematoxylin (two right panels). Dark and light blue bars indicate the 5.8S rRNA immunoreactivity values found in basal and suprabasal epithelial layers, respectively. Scale bar, 10 µm.

**(B)** Quantitation of the 5.8S rRNA immunoreactivity obtained in panel (A). ***, $P < 0.0001$ of test samples versus their respective layer control (ANOVA and Tukey's HSD tests, $n = 3$ independent experiments). Data represent the mean ± SEM. Source data for this figure are provided as a Source Data file.

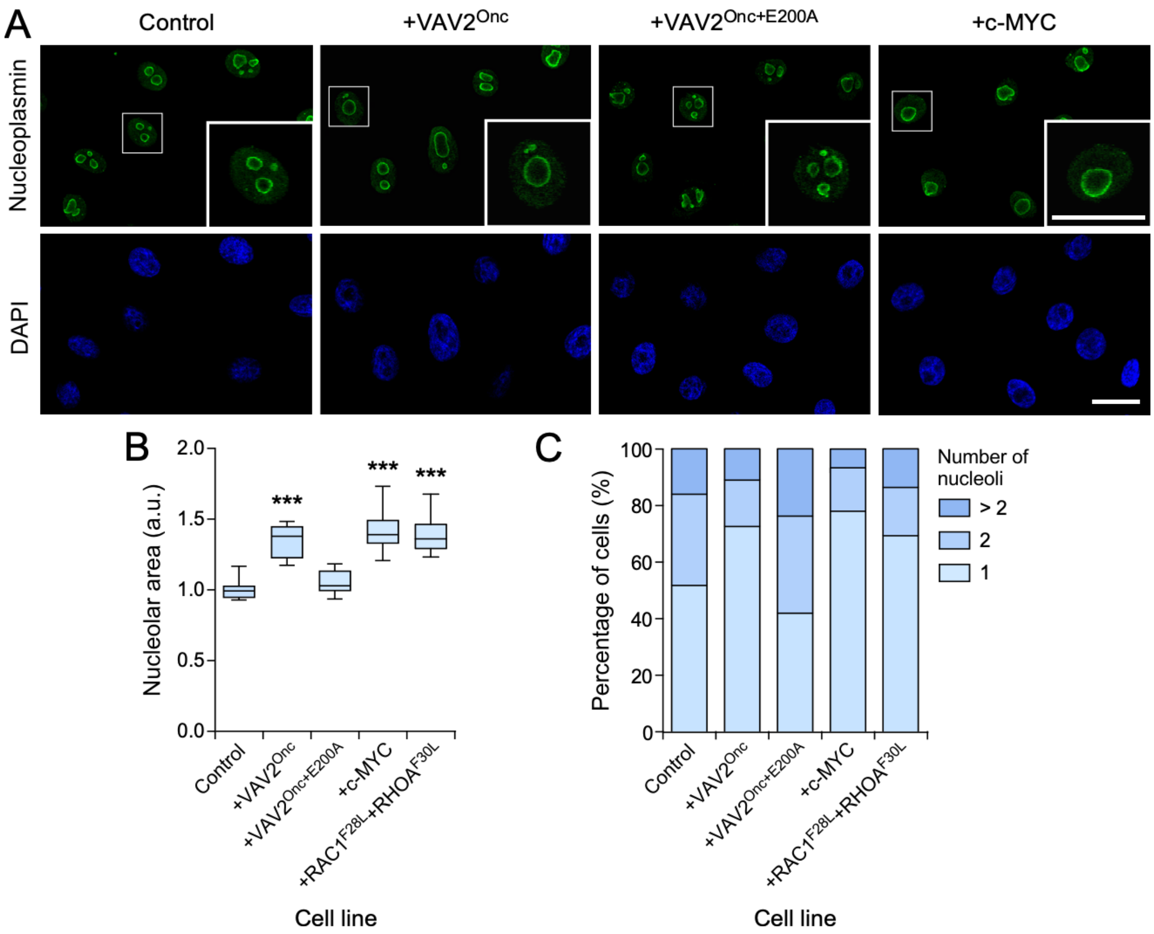


**SUPPLEMENTARY FIGURE 3. Activation of the VAV2 pathway promotes changes in nucleolar morphology**

**(A)** Representative confocal microscopy showing the type of nucleoli found in human keratinocytes expressing the indicated proteins (top). Nucleoli and nuclei were detected using immunostaining with the nucleolar protein nucleoplasmin (top panels) and DAPI staining (lower panels), respectively. Scale bar, 20 µm. In the top panels, the squares indicate the areas that are shown enlarged in the right bottom corner of each image.

**(B and C)** Quantitation of the area (B) and number (C) of nucleoli found in keratinocyte samples analyzed in (A). ***, $P < 0.001$ (ANOVA and Dunnett's multiple comparison tests, $n = 3$ independent experiments). Data represent the mean ± SEM. Source data for this figure are provided as a Source Data file.

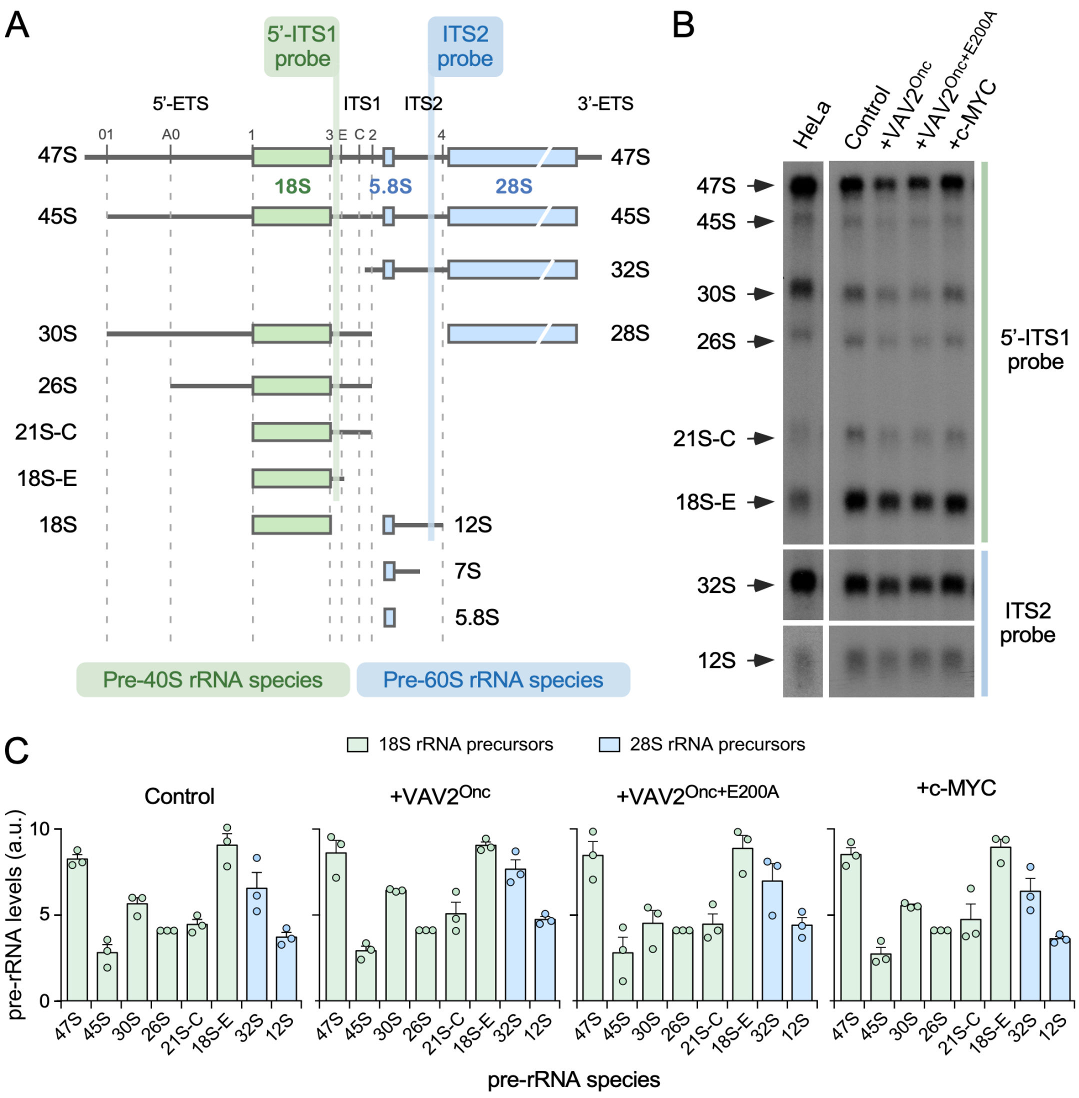


**SUPPLEMENTARY FIGURE 4. VAV2$^{Onc}$ does not induce changes in pre-rRNA processing in human keratinocytes**

**(A)** Scheme depicting the pre-rRNA processing pathway in human cells. Pre-rRNA species belonging to the small (left, green) and large (right, blue) ribosome subunits can be identified by Northern blot using the 5′-ITS1 and the ITS2 probes (top), respectively. The size of the pre-rRNA species is indicated on the right and on the left of the figure. ETS, external pre-rRNA segment; ITS, internal pre-rRNA segment.

**(B)** Representative Northern blot analysis using total RNAs from HeLa cells (left column) and indicated human keratinocytes (rest of lanes) to detect the abundance of the pre-rRNA species that are recognized by the 5'-ITS and ITS2 probes (right). The size of the pre-rRNA species is indicated on the left.

**(C)** Quantitation of the relative abundance of the interrogated pre-RNA species in the keratinocyte cell lines analyzed in (B). Normalization was performed relative to the abundance of the 47S pre-RNA. Not statistically significant differences were found between the indicated samples and controls (ANOVA and Dunnett's multiple comparison test, $n = 3$ independent experiments). Data represent the mean ± SEM. Source data for this figure are provided as a Source Data file.

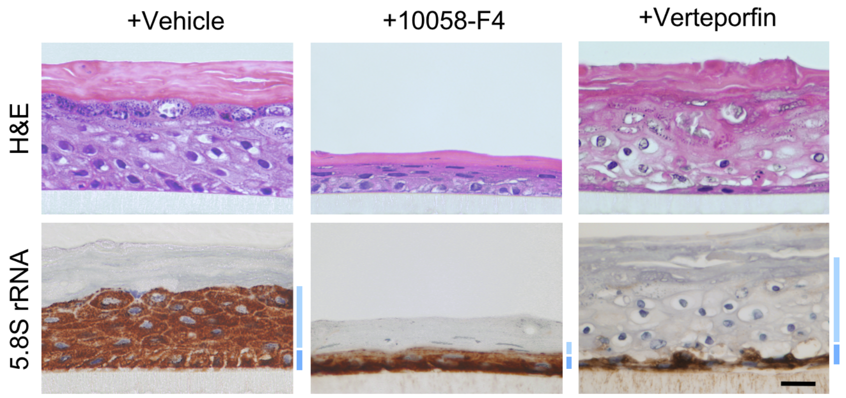


**SUPPLEMENTARY FIGURE 5. Downstream VAV2 GTPases also promote increased synthesis of the pre-rRNA in a MYC- and YAP/TAZ-dependent manner**

Representative confocal microscopy images of RAC1$^{F28L}$+RHOA$^{F30L}$-expressing human keratinocytes treated with the indicated inhibitors (top) that were stained with either hematoxylin-eosin (top panels) or with an antibody to the 5.8S rRNA plus hematoxylin (bottom panels). Dark and light blue bars indicate the basal and suprabasal epithelial layers, respectively (right). Scale bar, 10 µm. The quantitation of 5.8S rRNA immunoreactivity found in this and two additional experiments is shown in **Figure 3B** (right).
